# Cerebellar Activity in Hemi-Parkinsonian Rats during Volitional Gait and Freezing

**DOI:** 10.1101/2023.02.28.530475

**Authors:** Valerie DeAngelo, Arianna Gehan, Siya Paliwal, Katherine Ho, Justin D Hilliard, Chia-Han Chiang, Jonathan Viventi, George C McConnell

## Abstract

Parkinson’s disease is a neurodegenerative disease characterized by gait dysfunction in the advanced stages of the disease. The unilateral 6-OHDA toxin-induced model is the most studied animal model of Parkinson’s disease, which reproduces gait dysfunction after greater than 68% dopamine (DA) loss in the substantia nigra pars compacta (SNc). The extent to which the neural activity in hemi-parkinsonian rats correlates to gait dysfunction and DAergic cell loss is not clear. In this paper we report the effects of unilateral DA depletion on cerebellar vermis activity using micro-electrocorticography (μECoG) during walking and freezing on a runway. Gait and neural activity were measured in 6-OHDA lesioned and sham lesioned rats at 14d, 21d, and 28d after infusion of 6-OHDA or control vehicle into the medial forebrain bundle (MFB) (*n*=20). Gait deficits in 6-OHDA rats were different from sham rats at 14d (*p*<0.05). Gait deficits in 6-OHDA rats improved at 21d and 28d except for run speed, which decreased at 28d (*p*=0.018). No differences in gait deficits were observed in sham lesioned rats at any time points. Hemiparkinsonian rats showed hyperactivity in the cerebellar vermis at 21d (*p*<0.05), but not at 14d and 28d, and the activity was reduced during freezing epochs in lobules VIa, VIb, and VIc (*p*<0.05). These results suggest that DAergic cell loss causes pathological cerebellar activity at 21d postlesion and suggests that compensatory mechanisms from the intact hemisphere contribute to normalized cerebellar activity at 28d. The decrease in cerebellar oscillatory activity during freezing may be indicative of neurological changes during freezing of gait in Parkinson’s disease patients making this region a potential location for biomarker detection. Although the unilateral 6-OHDA model presents gait deficits that parallel clinical presentations of Parkinson’s disease, further studies in animal models of bilateral DA loss are needed to understand the role of the cerebellar vermis in Parkinson’s disease.

## Introduction

Parkinson’s disease is a progressive neurodegenerative disorder that causes symptoms of gait dysfunction, including freezing of gait (FOG). Although the exact mechanism is unclear, gait deficits typically appear with substantia pars compacta (SNc) dopamine (DA) loss exceeding 40% and become pronounced after 68% loss.^1–6^ Symptoms present as shuffled gait, decreased stride length, and decreased velocity.^6^ FOG is the most difficult symptom to treat and is defined as the episodic absence or marked reduction of forward progression of gait despite the intention to walk.^7–8^

Studies have found that symptoms of Parkinsonian gait are consistently replicated in the unilateral 6-hydroxydopamine (6-OHDA) model of Parkinson’s disease following 68% DA loss.^6–8^ Sufficient DA loss for gait dysfunction occurs as early as 7-days post 6-OHDA infusion into the medial forebrain bundle (MFB) and the gait dysfunction is dose dependent.^9^ Zhou et al.^10^ suggested that an increased duration of stance time may indicate FOG and gait hesitation. The similarities between the acute 6-OHDA gait deficits and clinical presentations in advanced Parkinson’s disease support the assertions that this model has good face validity to further our understanding of the circuit mechanisms involved in Parkinsonian gait.^11^

The cerebellum - a brain region important for motor planning, control, and execution - is known to exhibit aberrant activity in patients with Parkinson’s disease.^12–13,15–18^ Cerebellar vermis (CBLv) contributions to gait dysfunction in Parkinson’s disease has been linked to degeneration in the pedunculopontine nucleus (PPN) and basal ganglia (BG).^1,14,44^ To our knowledge, the current understanding of CBLv changes in Parkinson’s disease are limited to results from imaging studies which measure activity indirectly by coupling activity to increases in blood flow^20^ and EEG recordings, which have limited spatial resolution.^19^

Hanakawa et al. found that the CBLv is hyperactive in Parkinson’s disease patients experiencing gait dysfunction. Most studies that have measured CBLv hyperactivity in Parkinson’s disease were performed indirectly using imaging methods. These methods typically measure activity before, after, or in the absence of gait.^12–13,15–18^

A more recent electrophysiological study by Bosch et al. using EEG found CBLv activity was attenuated during cognitive processing and lower limb performance in Parkinson’s disease subjects compared to healthy controls.^19^ To our knowledge this is one of the few studies to measure the electrophysiology of the CBLv.

We measured the oscillatory activity of CBLv lobules VIa, VIb, VIc, and VII in 6-OHDA rats using micro-electrocorticographic (μECoG) arrays.^21^ The μECoG arrays enabled high spatial resolution electrophysiological recordings from CBLv during freezing and continuous gait and the ability to discriminate individual lobules with the CBLv.^21^ Gait and power spectral analysis performed 14-, 21-, and 28-days after 6-OHDA infusion allowed us to directly measure the effects of DA depletion in the SNc on gait and cerebellar activity. Our results provide insight into the face validity of the unilateral 6-OHDA model for gait dysfunction in Parkinson’s disease and the underlying neural activity in the CBLv.

## Material and Methods

### Animals and housing

Adult male Long-Evans hooded rats (*n* = 25) were purchased from Charles River Laboratories. All rats were between 4 months and 5 months of age and were singly housed (postsurgery) in standard cages with free access to food and water. Study protocols were reviewed and approved by the Stevens Institute of Technology Institutional Animal Care and Use Committee.

### Experimental design

6-OHDA infusion in the MFB causes degeneration of DAergic neurons in the SNc. To measure the effect of DA loss on cerebellar activity, three groups of rats were lesioned with 6-OHDA: 14-day (*n* = 7), 21-day (*n* = 7), and 6-OHDA (*n* = 6). The 14-day and 21-day groups were perfused 14- and 21-days post-lesion, respectively. Rats in the 6-OHDA group were chosen at random and implanted with a μECoG array to longitudinally record CBLv activity during gait. The control group (*n* = 5), chosen at random and labeled sham, was infused with sterile saline and implanted with a μECoG array. 6-OHDA and sham rats were perfused at 28-days post-surgery. Neural and gait analyses were performed unblinded while immunohistochemistry and cell counting were performed blinded.

### Unilateral 6-OHDA lesion and chronic μECoG implantation

Sterile stereotaxic surgery was conducted under 4% sevoflurane anesthesia using coordinates from a rat brain atlas.^22^ Rats were placed in a stereotaxic frame and a craniotomy was made over the CBLv (AP −9.5mm to −14.5mm, ML ±4mm) and the MFB (AP −1.8mm, ML + 2.0mm, DV −8.6mm relative to bregma). Eight stainless steel screws were anchored to the skull, including two screws placed over the visual cortex which served as reference and ground. Thirty minutes prior to infusion rats were pretreated with intraperitoneal (IP) injections of 50mg/kg pargyline (Sigma-Aldrich) and 5mg/kg desipramine (Sigma-Aldrich) to inhibit monoamine oxidase and protect non-adrenergic neurons, respectively.^23^

Immediately before infusion, 6-OHDA hydrobromide (Tocris Bioscience) was dissolved in ice-cold 0.02% ascorbic acid dissolved in saline to a concentration of 4 μg/μl. 4 μl of 6-OHDA was infused into the MFB of the right hemisphere at a rate of 1 μl/min for a total of 16μg. Sham rats did not receive desipramine/pargyline and were infused with 4 μl of sterile saline in lieu of 6- OHDA. Immediately following infusion, a μECoG array, arranged in a 8×8 grid of 61 total electrodes with 230 μm diameter electrodes spaced 420μm (Fig. 1A), was implanted over CBLv lobules VIa (AP −11.205 mm to −11.855mm, ML ±1.5mm), VIb (AP −12.045 mm to −12.7mm, ML ±1.5mm), VIc (AP −12.88 mm to −13.21mm, ML ±1.3mm), and VII (AP −13.4mm to −14.05mm, ML ±1.5mm) in 6-OHDA and sham rats (Fig 1B-D). Dental acrylic was used to secure the μECoG array to stainless steel screws (Fig. 1E). Sterile Covidien Vaseline petroleum jelly was placed over the craniotomy to create a barrier between the dental acrylic head cap and the brain surface. Electrode location was confirmed after perfusion using a vernier caliper.

**Figure 1:**
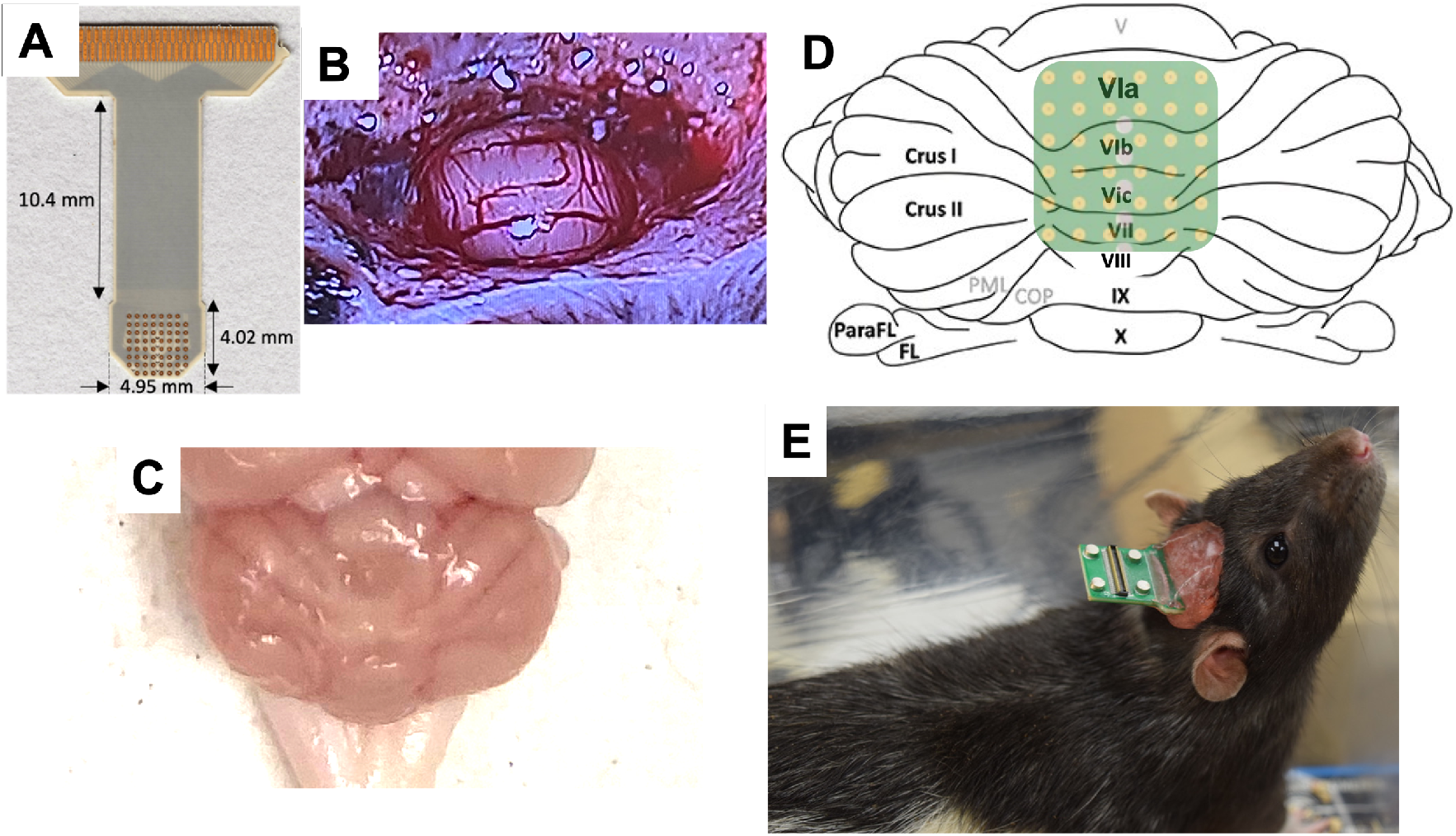
Chronic μECoG implantation. (A) μECoG array arranged in an 8×8 grid with 230μm diameter spaced 420μm apart. (B) Craniotomy created over cerebellar vermis (C) lobules VIa through VII. (D) Approximate location of the μECoG array when chronically implanted. (E) Finished head cap.

### Methamphetamine-induced circling

Seven days post-lesion, rats from the 6-OHDA, 14-day, and 21-day groups were administered 1.6mg/kg of methamphetamine through intraperitoneal injection and placed in a dark circling chamber for 1.5 hours. An infrared camera captured the rat’s activity. The lesion was considered successful if the rat circled at a minimum rate of 3 turns per minute indicative of >80% dopamine loss.^23^ Rats that did not have a successful lesion based on this criteria were excluded from the remainder of this study.

### Gait analysis

Gait assessment was performed on the 6-OHDA (*n* = *5*) and sham (*n* = *5*) rats using a runway system for gait analysis (CSI-G-RWY; CleverSys Inc., Reston, VA). As the rat voluntarily moved across the runway, comprised of a long side-lit glass plate, each footprint was lit up and recorded by a camera mounted under the glass. Prior to assessment, each rat was trained daily for three consecutive runs until they walked across the apparatus without hesitation. Each rat was then evaluated at 14-days, 21-days, and 28-days post-lesion. Data from three consecutive runs were averaged for each rat at each timepoint. Each video was analyzed using the gait analysis software (GaitScan; CleverSys Inc., Reston, VA). After 5 minutes, if the 6-OHDA subject would not walk on the runway, they were placed back in their home cage for 3 minutes and an attempt was made again.

Analysis was performed for periods of walking (continuous gait consisting of at least three consecutive steps) and freezing. Freezing events were defined as cessation of continuous gait with at least 3 paws in contact with the glass for greater than 500ms. Analysis was performed on swing time (in ms; time in which paw is in the air), stance time (in ms; time in which all paws are detected on the glass), stride time (in ms; stance time plus swing time), stride length (in mm; distance the paw traversed from start of previous stance to beginning of next stance) and run speed (instantaneous speed over a running distance).^24^

### Neural recordings

Local field potential (LFP) recordings were taken during volitional gait on the runway using the Open Ephys (open-source electrophysiology) system (Fig. 2A and B).^2^ The runway was modified to synchronize LFP recordings with gait. Two beam break sensors were used in conjunction with an Arduino to control the timing of the LFP recording period. As the rat left its start box and entered the field of view, the first sensor was broken, a TTL pulse was sent to Open Ephys to start recording, and an LED was turned on. LFPs were continuously recorded as the rat traversed the runway. After the rat exited the runway and entered the exit box, a second sensor was broken sending a TTL pulse to stop the recordings and turn off the LED. The LED served as a visual indicator for the start and end of LFP recordings during gait analysis. Data from three consecutive runs were stored for each rat for further analysis.

**Figure 2:**
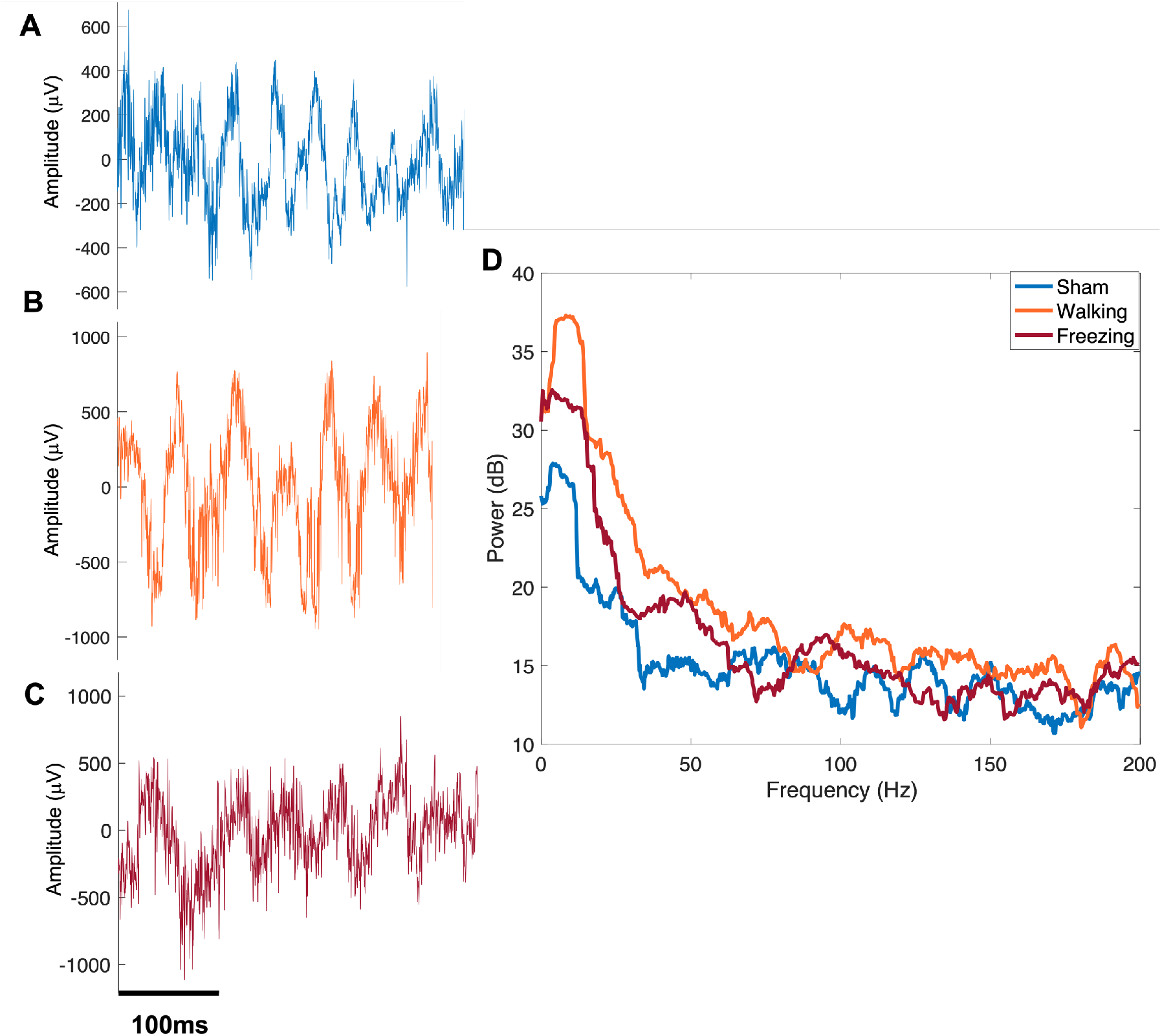
Sample neural recordings. (A) Sample LFPs recorded from cerebellar vermis lobule VIa of sham control during walking and 6-OHDA during (B) walking and (C) freezing. (D) Sample power spectra (dB) from LFPs measured in sham (blue), 6-OHDA during walking (orange) and 6- OHDA during a bout of freezing (red) during volitional gait.

### Neural Data Analysis

Continuous LFP recordings were exported and analyzed using MATLAB (Mathworks, Natick, MA). First, LFP timing was synchronized with the LED ON and OFF video frames to crop recordings based on walking and freezing for 6-OHDA rats as sham rats did not freeze on the runway. Next, LFP recordings were separated into epochs of walking and freezing based on the runway videos. LFP recordings during the first ¾ of the length of the runway were analyzed, because most rats (sham and 6-OHDA) stopped at the end of the runway prior to exiting into their home cage. Multitaper methods of spectral estimation were used to quantify changes in CBLv oscillatory activity between 6-OHDA (Fig. 2B-C) (walking and freezing) and sham rats (Fig. 2A) (walking) in each lobule (Chronux version 2.00) (Fig. 2D). The LFP spectrum was estimated on a 2 second window with 10 Hz resolution using 19 Slepian data tapers.^23^ The mean power in dB was measured within the following frequency bands: theta (4-8Hz), alpha (8-13Hz), low beta (13-20Hz), high beta (21-30Hz), low gamma (30-55Hz), high gamma (65-80Hz) and fast frequency (80-200Hz). Frequencies between 55Hz and 65Hz were omitted to reduce effects of 60Hz noise.^25^

### Immunohistochemistry

Fourteen days, 21-days, and 28-days post-lesion rats were deeply anesthetized and transcardially perfused with PBS followed by 10% formalin. The brain was removed and postfixed overnight (4°C) in formalin then placed in a 30% sucrose (4°C) solution until it sank. A green tissue marking dye was applied to the left-posterior hemisphere to demarcate the orientation of the brain sections. The brains were cryoprotected with Tissue-Tek optimal cutting temperature (O.C.T.) compound and 40μm serial coronal sections were cut using a cryostat (CryoStar NX50) equally spaced through the substantia nigra pars compacta (SNc). Immunohistochemistry was performed in the SNc with anti-tyrosine hydroxylase (TH) antibody to measure DAergic neuron loss.^24^ After three rinses in phosphate buffered saline (PBS), sections were blocked for 1 hour at 4°C in blocking solution containing PBS, normal goat serum (NGS), and 10% Triton-X. The sections were then incubated in anti-TH monoclonal rabbit IgG primary antibody (1:2000 overnight at 4°C; Sigma-Aldrich) in solution with PBS and normal goat serum (NGS). Next, the sections were incubated with goat anti-rabbit IgG Alexa 488 secondary antibody (1:500 for 2 hours at 4°C; Invitrogen by ThermoFisher Scientific) in solution with PBS, NGS, and 10% Triton-X. Sections were mounted in FluoroMount-G and imaged using the Keyence BZ-X700 series microscope (Supplementary Fig. 1).

### Cell counting

Automated cell counting was performed using ImageJ.^24,26–27^ DAergic degeneration was determined in the SNc by counting the number of TH-positive cells of each rat (Supplementary Fig. 1). Each 10x image was converted from RGB to 16-bit greyscale. The threshold was set to a pixel intensity of 30 (arbitrary units) and cells were analyzed in a region of interest (ROI) restricted to a 705.9 mm (w) x 564.7 mm (h) rectangle on either side of the midline in the SNc,

The percentage of DA loss of the lesioned side normalized to the non-lesioned side was measured using the following equation^23^:

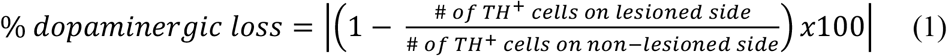

### Statistical analysis

Statistical analysis was performed using IBM SPSS Statistics for Mac (IBM Corp., Armonk, N). Means were compared using a two-way mixed ANOVA with time and type as factors for gait and LFP power comparisons between 6-OHDA and sham rats. For significant interactions univariate analysis and one-way repeated measures ANOVA was used to determine significance between types at each time point and between timepoints for each type, respectively. Percent DA loss of the 14-day, 21-day, 6-OHDA (28-day), and sham rats (control) were compared using a oneway ANOVA. Sidak post hoc test was used for all analyses. Results were considered statistically significant at *p* ≤ 0.05.

## Results

### Methamphetamine-induced circling

A total of 20 rats were administered 6-OHDA in this study (14-day (*n* = 7), 21-day (*n* = 7), and 28 day (*n* = 6)). Of these 20 rats, 5 lesions did not meet the minimum of 3 turns per minute over the period of 1.5 hours post-6-OHDA-injection and excluded from further analysis (data not shown).

### Gait analysis

Gait dysfunction was observed across all measurements between 6-OHDA (*n* = 5) and sham control rats (*n* = 4). Gait analysis for one of the sham control rats could not be performed due to data corruption of the video files. 6-OHDA rats had periods of freezing at the beginning, middle, and end of the runway at each timepoint while the sham rats only paused at the end when entering their home cage. This was removed from analysis as it was not deemed as a freezing episode. Start hesitation was also observed at each time point. During multiple runs, 6-OHDA continuously circled at the start box which resulted in no forward progression of gait. Placing the rat back in their home cage for 3 minutes and restarting the trial typically resulted in a successful run where the rat was able to traverse down the runway allowing gait measurement.

Stride time, stance time, and swing time were compared between 6-OHDA rats and sham controls. 6-OHDA rats experienced gait deficits when compared to age matched sham controls only at 14-days post-lesion. No statistically significant interaction between type and time was found for stance time, swing time, and stride time. Significant differences were found for the main effect of type for stride time (*F*(1,7) = 17.431, *p* = 0.004) and stance time (*F*(1,7) = 20.637, *p* = 0.003), but not swing time (*F*(1,7) = 3.462, *p* = 0.105). 6-OHDA rats had greater stride time and stance time when compared to the sham control. Simple effects analysis revealed significant differences between 6-OHDA and sham at 14 days but not at 21 days or 28 days for stride time (*F*(1,7) =33.820, *p* < 0.001), stance time (*F*(1,7) = 41.366, *p* <0.001), and swing time (*F*(1,7) = 5.820, *p* = 0.047). No significant main effect of time was found across temporal gait measures. Overall, stance time, stride time and swing time were greater in 6-OHDA lesioned rats than sham lesioned at all timepoints (Fig. 3).

**Figure 3:**
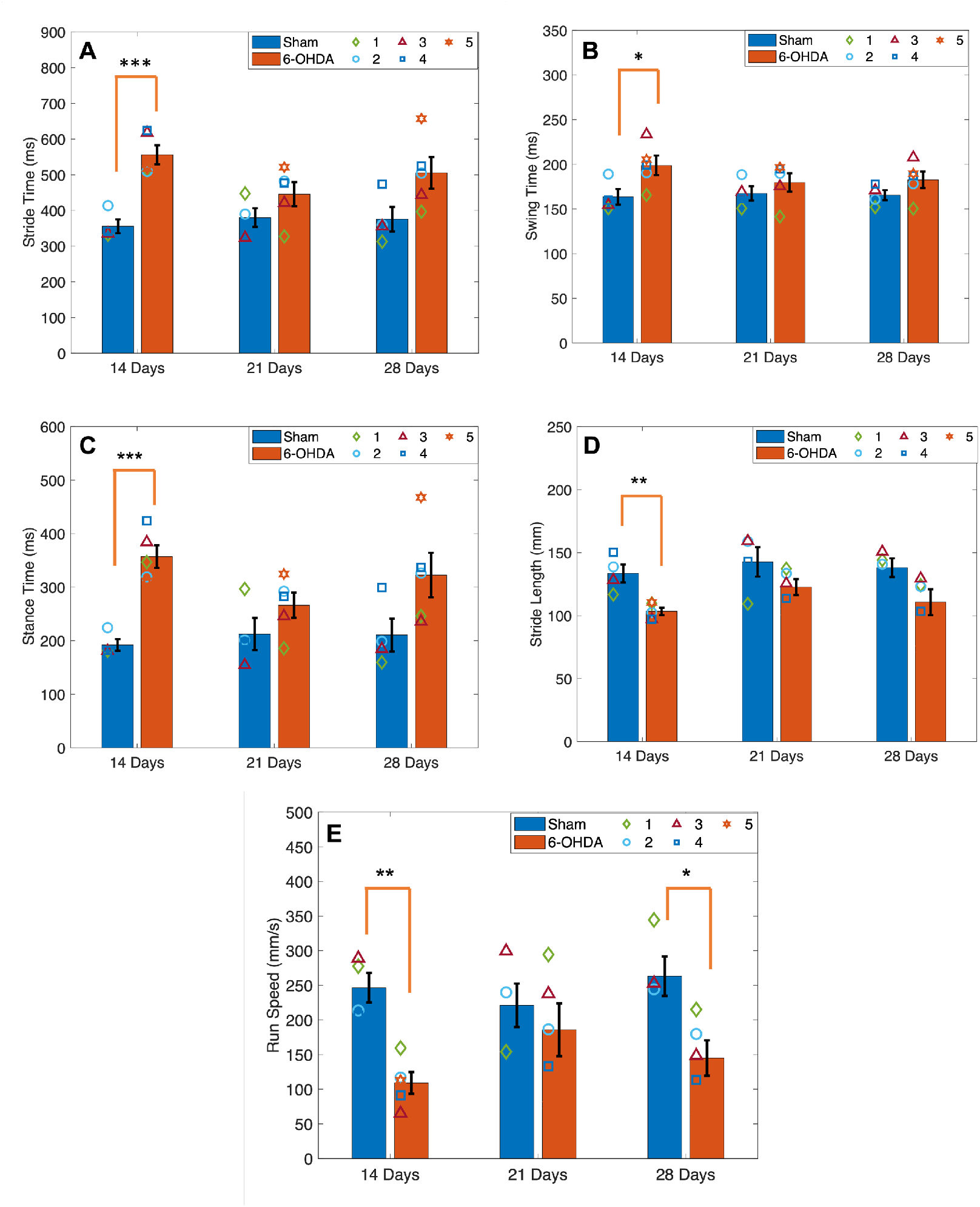
Gait analysis. (A) Stride time, (B) swing time, and (C) stance time were significantly greater in 6-OHDA lesioned rats compared to sham at 14 days but not at 21 and 28 days. (D) Stride length was significantly greater in 6-OHDA rats at 14 days compared to sham controls. (E) 6- OHDA had a significantly slower run speed than sham controls at both 14 days and 28 days, but not 21 days. Symbols represent the mean at each timepoint in 6-OHDA rats (n = 5) and sham control rats (n = 4). * indicates p < 0.05, ** indicates p < 0.01, and *** indicates p < 0.001. Bars indicate mean +/- SEM.

There was no significant two-way interaction between type and time for stride length (Fig. 3D). Main effect of type was significant (*F*(1,7) = 11.705, *p* = 0.011), while main effect for time was not. Simple main effects analysis found significant differences between 6-OHDA and sham rats only at 14-days (*F*(1,7) = 17.909, *p* = 0.004). Stride length was decreased at all timepoints in 6-OHDA rats compared to sham.

There was no significant two-way interaction between type and time on run speed. The main effect for type was statistically significant (*F*(1,7)=10.303, *p* = 0.015). Simple effects showed a significant reduction in speed at 14- (*F*(1,7) = 28.476, *p* = 0.001) and 28-days (*F*(1,7) = 9.563, *p* = 0.018) in 6-OHDA compared to sham. (Fig. 3E). No significant main effect was found for time.

### Mean power

LFP average logarithmic power was calculated in CBLv lobules VIa, VIb, VIc and VII during periods of walking and freezing in the delta, theta, alpha, low beta, high beta, low gamma, high gamma, and fast frequency bands. Activity during walking was compared between 6-OHDA and sham rats at 14-, 21-, and 28-days after surgery. Mean power during freezing was compared to mean power during periods of walking in 6-OHDA lesioned rats at all time points. Vermis lobules of 6-OHDA rats were hyperactive at 14 and 21 days when compared to sham rats in all frequency bands in all lobules. This activity normalized to levels of the sham group at 28-days.

### Walking

No statistically significant interaction between type and time was found for any of the frequency bands in all lobules during walking. The lobule VIa main effect type was significant in the high beta (*F*(1,7) = 6.277, *p* = 0.041) and low gamma (*F*(1,7) = 7.794, *p* = 0.027) bands (Fig. 4). Simple effects analysis revealed significant hyperactivity in 6- OHDA lesioned rats at 21 days in the low beta (*F*(1,7) = 7.2, *p* = 0.031), high beta (*F*(1,7) = 10.204, *p* = 0.015), low gamma (*F*(1,7) = 14.769, *p* = 0.006), and high gamma (*F*(1,7) = 7.105, *p* = 0.032) bands compared to sham controls.

**Figure 4:**
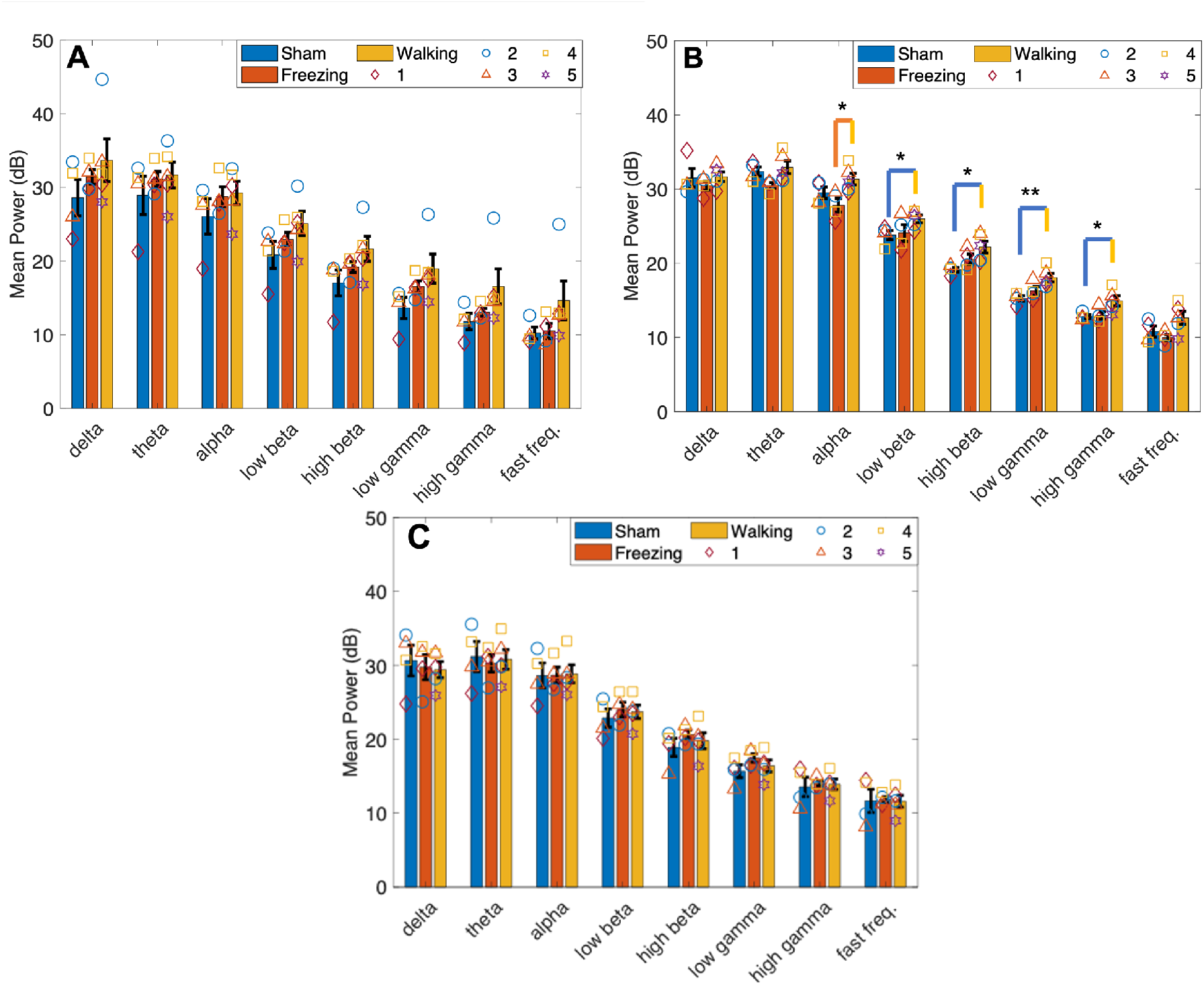
Mean power in lobule VIa. (A) Mean power measured in sham rats during walking and 6-OHDA rats during walking and freezing 14 days post-lesion. No significant differences were found between type. (B) Mean power measured in sham and 6-OHDA rats at 21 days post-lesion. Hyperactivity was measured in 6-OHDA rats during walking in the low beta, high beta, low gamma, and high gamma frequency bands compared to sham. A significant difference was measured between walking and freezing in 6-OHDA rats in the alpha band (C). At 28 days no significant differences in power were measured. Symbols represent the mean at each timepoint in (*n* = 5) 6-OHDA rats and (*n* = 4) sham control rats. * indicates p < 0.05; ** indicates *p* < 0.01. Bars indicate mean +/- SEM.

Main effect time was significant in lobule VIb in the alpha (*F*(2,14) = 4.932, *p* = 0.024 and low beta (*F*(2,14) = 4.102, *p* = 0.040) bands (Fig. 5). Pairwise analysis found power was significantly higher at 21-days post-lesion compared to 28-days in both frequency bands (alpha: *p* = 0.042, low beta: *p* = 0.046). Post hoc analysis of lobule VIb power using a one-way repeated measures ANOVA found a significant decrease from 21- to 28-days in the alpha (*p* = 0.042) and low gamma (*p* = 0.024) bands only in the 6-OHDA lesioned group (Fig. 8D). No significant main effect for time or type was found in lobules VIc (Fig. 6) and VII (Fig. 7).

**Figure 5:**
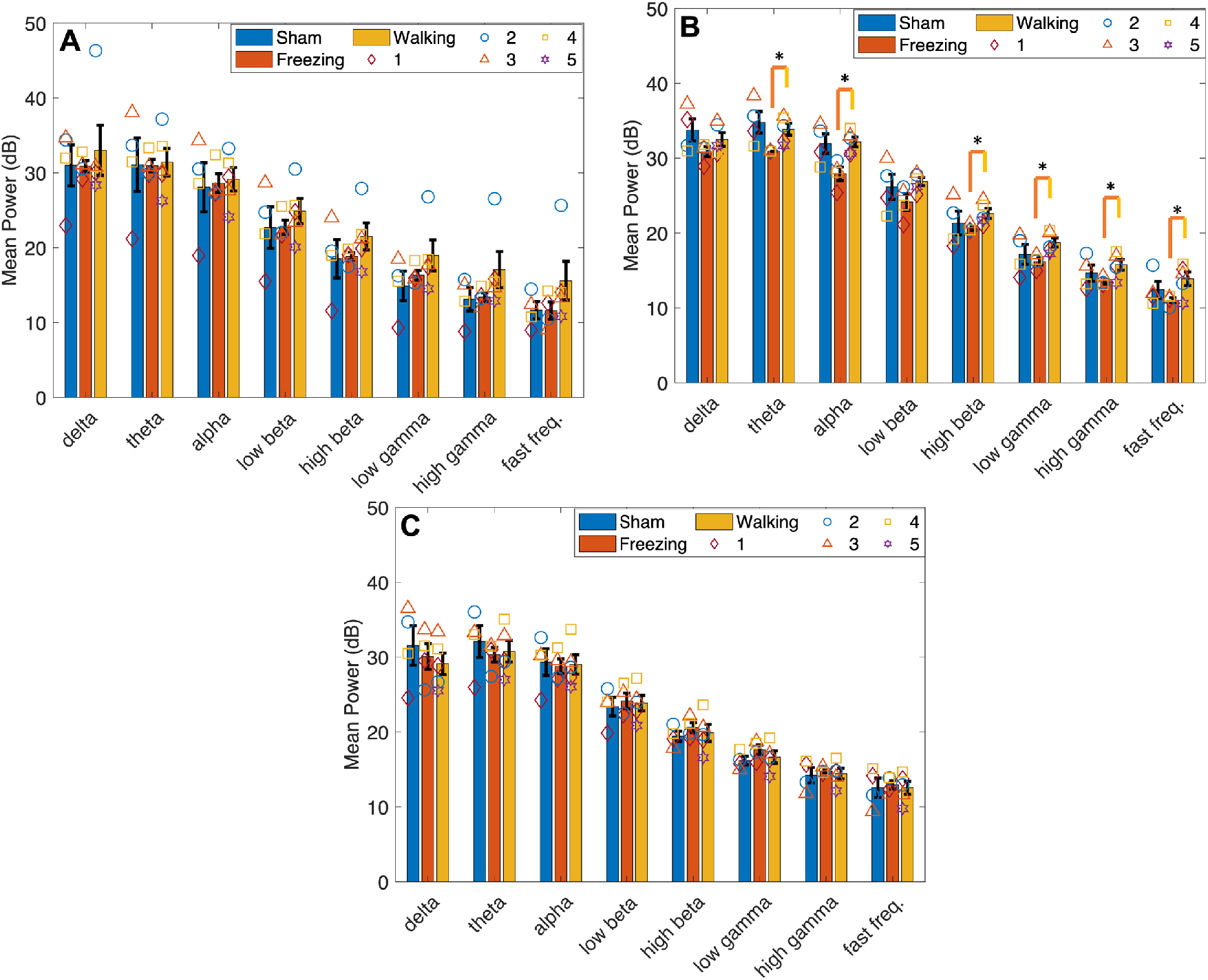
Mean power in lobule VIb. (A) No significant differences were found in mean power measured in sham rats during walking and 6-OHDA rats during walking and freezing 14-days post-lesion. (B) Mean power measured in sham and 6-OHDA rats at 21-days post-lesion. Hypoactivity was measured in 6-OHDA rats during freezing in the theta, alpha, high beta, low gamma, and high gamma, and fast frequency bands compared to walking. No significant differences were measured during walking between power in 6-OHDA and sham rats. (C) At 28-days no significant differences in power were measured. Power was significantly higher at 21-days post-lesion compared to 28-days in 6-OHDA rats in the alpha and low gamma bands during walking. These associations can also be seen in figure 8C. Symbols represent the mean at each timepoint in (*n* = 5) 6-OHDA rats and (*n* = 4) sham control rats. * indicates p < 0.05. Bars indicate mean +/- SEM.

**Figure 6:**
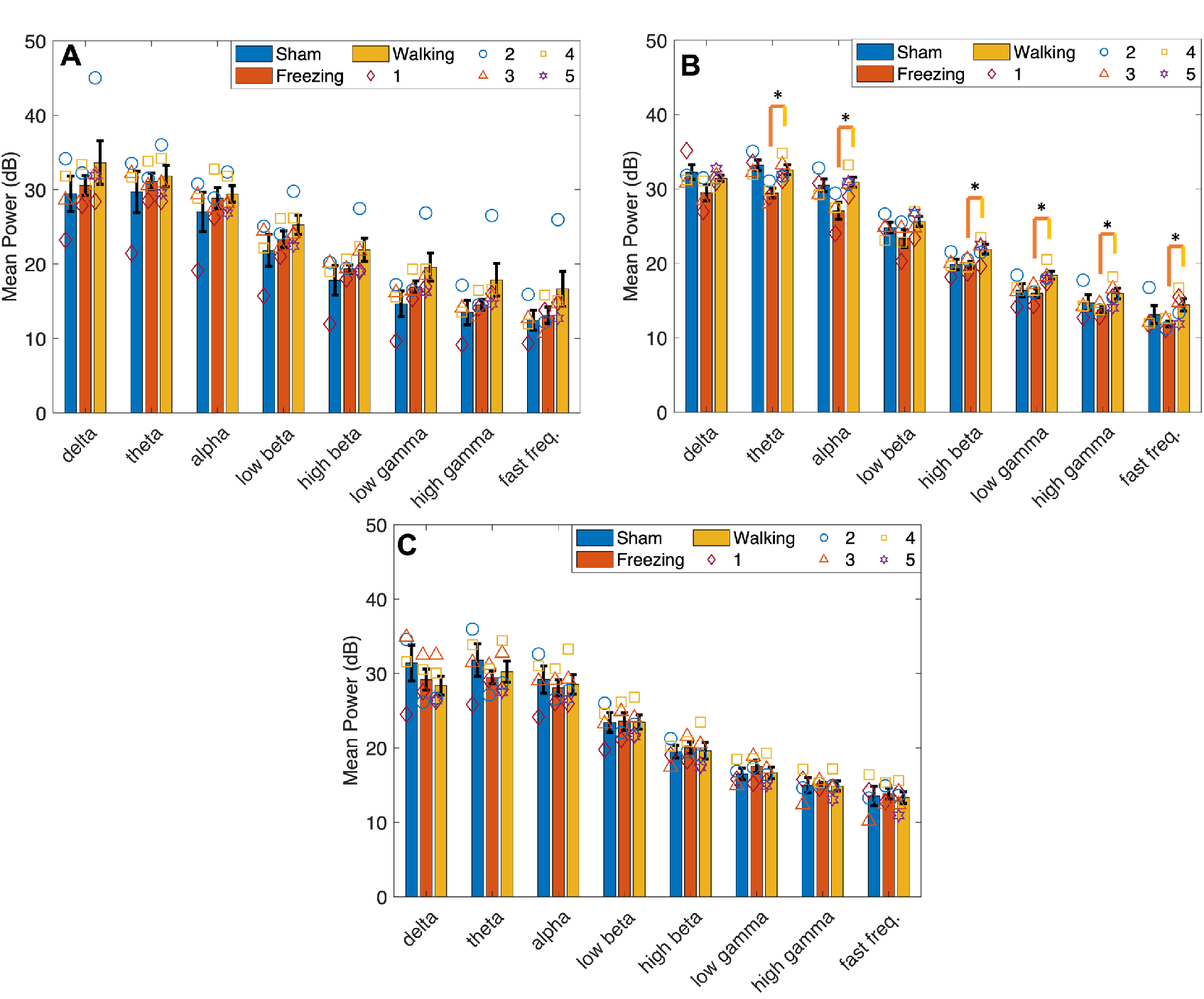
Mean power in lobule VIc. (A) No significant differences were found in mean power measured in sham rats during walking and 6-OHDA rats during walking and freezing 14-days after lesion surgery. (B) Mean power measured in sham and 6-OHDA rats at 21-days post-lesion. Hypoactivity was measured in 6-OHDA rats during freezing in the theta, alpha, high beta, low gamma, and high gamma, and fast frequency bands compared to walking. No significant differences were measured during walking between power in 6-OHDA and sham rats. (C) No significant differences in power were measured 28-days post-lesion. Symbols represent the mean at each timepoint in (*n* = 5) 6-OHDA rats and (*n* = 4) sham control rats. * indicates p < 0.05. Bars indicate mean +/- SEM.

**Figure 7:**
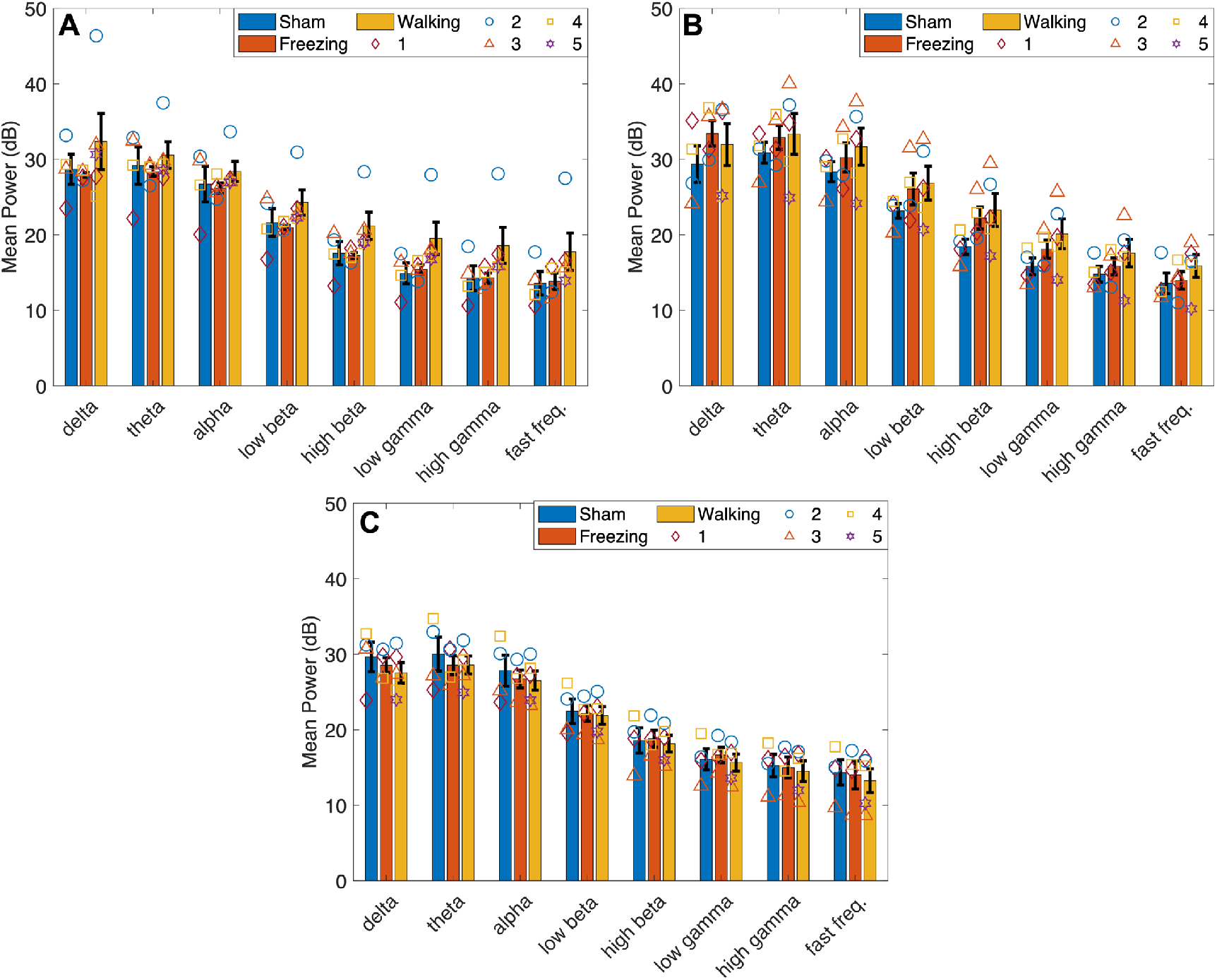
Mean power in lobule VII. (A) No significant differences were found in mean power measured in sham rats during walking and 6-OHDA rats during walking and freezing 14-, (B) 21 and (C) 28-days post-lesion surgery. A significant decrease in power was measured in the theta, alpha, low beta, and high beta band from 14- to 21-days (A-B) and an increase in power in the delta, theta, and high beta bands from 21- to 28-days (B-C). These associations can also be seen in figure 8E. Symbols represent the mean at each timepoint in (*n* = 5) 6-OHDA rats and (*n* = 4) sham control rats. Bars indicate mean +/- SEM.

**Figure 8:**
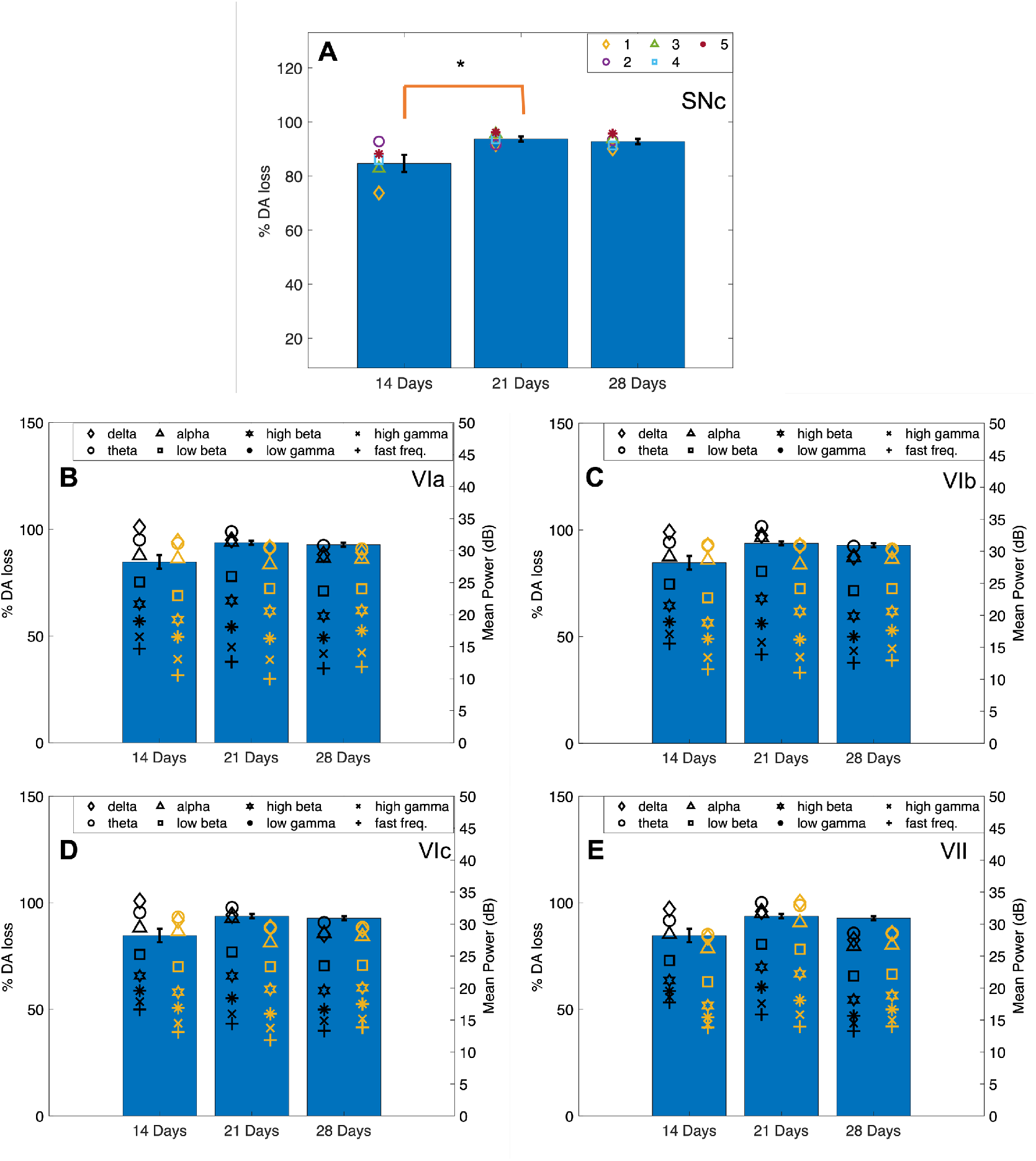
Cell counts and LFP activity in cerebellar vermis. (A) Percent dopamine loss increased in the SNc from 14 days and 21-days then remained constant from 21-days to 28-days post-lesion. Each point represents the mean percent dopamine loss in 6-OHDA rats between 14- (*n* = 5), 21- (n=5), and 28-days (n=5). * indicates p < 0.05. (B-E) Plots displaying the effect of dopamine loss in the SNc on cerebellar vermis power during walking and freezing in each lobule. Each point represents the mean power in each frequency band. Black symbols indicate LFP power during walking and yellow symbols represent LFP power during freezing at 14-, 21-, and 28-days. Bars indicate mean +/- SEM.

### Freezing

Fourteen days after 6-OHDA lesioning power during walking was largest compared to freezing and sham. Power during freezing was reduced to values greater than (lobules VIa and VIb) (Fig. 4 and 5) or equal to (lobules VIc and VII) (Fig. 6 and 7) sham power.

At 21-days power during freezing exhibited hypoactivity in comparison to walking in both groups except in lobule VII, however, power during freezing was still larger than power measured by the sham group during walking. At 28-days, power during walking in 6- OHDA and sham and during freezing in 6-OHDA normalized to similar levels.

No statistically significant interaction between type and time was found for any of the frequency bands in all lobules during walking. No significant main effects for time or type were found in lobules VIa, VIb, and VIc. Main effect for time was found significant in the theta (*F*(2,14) = 5.232), *p* = 0.020), alpha (*F*(2,14) = 4.776), *p* = 0.026), low beta (*F*(2,14) = 5.362, *p* = 0.019), and high beta (*F*(2,14) = 5.235, *p* = 0.020) of lobule VII. Pairwise analysis revealed a significant decrease in power in the theta (*p* = 0.048), alpha (*p* = 0.046), low beta (*p* = 0.037) and high beta (*p* = 0.027) from 14 to 21 days, and a significant increase in power from 21-days to 28-days in the delta (*p* = 0.018), theta (*p* = 0.038), and high beta (*p* = 0.048) bands (Fig. 8F). Post hoc analysis found no further significant differences in power during walking or freezing individually in frequency bands between timepoints.

Simple effect univariate analysis at each timepoint found significant differences in power during walking and freezing in the alpha band of lobules VIa (*p* = 0.022), VIb (*p* = 0.006) and VIc (*p* = 0.018), the theta, high beta, low gamma, high gamma and fast frequency bands of lobules VIb (*p* = 0.014, *p* = 0.035, *p* = 0.017, *p* = 0.029, and *p* = 0.035) and VIc (*p* = 0.013, *p* = 0.018, *p* = 0.041, *p* = 0.019, *p* = 0.036, and *p* = 0.039) at 21-days post lesion. Power during freezing was significantly reduced from the power measured during walking at this timepoint.

### Cell Counting

Cell counting in the SNc resulted in a mean TH+ cells loss of 84.7% at 14-days, 93.7% at 21-days, and 92.7% loss at 28-days following the 6-OHDA lesion (Supplementary Fig. 2). Sham rats had a mean loss of 25% after 28 days. SNc DA loss was significant in 6-OHDA lesioned rats between 14 and 21 days (*p* = 0.026) but not 21- and 28-days (Fig. 8A).

Fig. 8B-E compares the TH+ cells DAergic cell loss in the SNc to mean power during walking and freezing at each time point. From 14- to 21-days mean LFP power in the delta, theta, and alpha frequency bands became more similar while low beta and high beta power increased. This was mimicked during freezing except in lobule VII in which power during freezing increased between timepoint. Power measured during freezing was lower than during walking at 14- and 21-days. At 28-days, however, this normalized where power during walking and freezing were more closely related across all lobules.

## Discussion

This study analyzed the effect of DA depletion in the SNc on gait dysfunction and CBLv activity in 6-OHDA rats. A μECoG array was placed over vermis lobules VI, VII, and VII due to their involvement in the regulation of complex movements and motor planning.^28–29^ These lobules receive signals by way of the fastigial nucleus (FN) from brain regions including the motor cortex, BG, and PPN, all of which are associated with gait dysfunction in Parkinson’s disease.^30^

All measurements were made at 14-days, 21-days, and 28-days following infusion with 6- OHDA and sterile saline. DAergic cell loss was over time was as follows: 84.7% at 14 days, 93.7% at 21 days, and 92.7% at 28 days. To ascertain differences in cerebellar activity over time it was important to use the same rats for gait and cerebellar recordings at all time points. Lesioning separate groups and perfusing them at 3 separate timepoints allowed us to be consistent with our measurements while still gaining an understanding of DA depletion over time. DA loss significantly increased from 14- to 21-days and plateaued from 21- to 28-days. This is consistent with what was reported by Hsieh et al.^31^, although the percent DA loss reported here was greater at 21-days (93.7% vs. 88.66%). The higher dosage of 6-OHDA (16μg vs. 8μg) we used may explain the increased severity of the DA lesion.

Gait analysis was performed during walking in 6-OHDA and sham lesioned rats at 14-, 21-, and 28-days post-lesion. 6-OHDA gait measurements at 14-days were significantly different from sham rats and consistent with measures previously reported.^8,10^ 6-OHDA rats experienced increased stance time, stride time, and swing time and decreased stride length and run speed when compared to sham controls. Gait measurements in sham rats remained consistent at all time points whereas 6-OHDA rats exhibited an improvement in gait deficits from 14- to 21-days, despite a significant increase in DAergic cell loss. Measurements at 21- and 28-days indicated deficits still existed but were no longer significantly different from sham controls, with the exception of run speed at 28-days.

6-OHDA rats moved more quickly at 21-days than at 14- and 28-days. Run speed was the only parameter to be significantly different from sham rats at 28-days. Improvements in gait may be a result of compensatory adjustments made by the non-lesioned side during gait. Hsieh et al.^31^ measured asymmetries between the lesioned and non-lesioned side in 6-OHDA rats over time. Gait deficits may not have been as severe as previously reported due to the exclusion of freezing events from our analysis of walking.

CBLv activity was recorded during volitional gait on a runway. The Arduino embedded system allowed us to synchronize LFP recordings to rat activity. The sensor was triggered when the rat entered the camera view, which allowed offline synchronization of the LFP activity with periods of continuous gait and freezing. Most 6-OHDA rats did not immediately engage in continuous gait upon being placed on the runway, despite the ability to do so during training. One reason for this may be start hesitation, a symptom seen clinically in FOG.^33^

A FOG episode initiated when approaching a doorway may be overcome by shifting attention to an object away from them thus distracting attention resulting in step length with the correct amplitude.^32^ If start hesitation lasted >5 minutes the rats were placed into their home cage for a 5-minute reset period. After the reset the rats were returned to the runways and, in most cases, engaged in active gait. We postulate that placing rats in their home cage created an attention shift, resetting normal function, which allowed them to initate gait when placed back on the runway. A limitation of this study was the inability to measure CBLv activity during this time. Future research may analyze cerebellar LFPs to see if any quantifiable changes in CBLv activity are seen during start hesitation between 6-OHDA rats and sham lesioned rats.

CBLv activity was hyperactive compared to sham at 14 days in all lobules and frequency bands, with the greatest differences in the higher frequency bands. This trend in activity persisted at 21-days. At this timepoint, vermis lobule VIa was significantly hyperactive compared to sham rats in the beta and gamma (low and high) bands. At 28-days, mean power in all frequency bands was reduced in all vermis lobules to similar values of sham rats.

Mean power was similar in lobules VIa, VIb, and VIc, across all time points, however, only VIa reached statistical significance between sham at 6-OHDA during walking at 21-days. Lobule VI has been shown to receive inputs to and from both the premotor and primary motor cortex^29,34^ and both lobules VI and VII are involved in cognition, emotion, and motor planning.^28–29^ Cortical activity is reduced in Parkinson’s disease.^1^ Hyperactivation of this lobule may be a response to impaired behavioral and motor planning due to a disconnection of the cortico-BG circuits, resulting in impaired gait. Although gait impairments were not statically significantly different at this timepoint, measures still showed deficits in the 6-OHDA rats compared to the controls.

Excessive beta activity (12-30Hz) is a prominent feature in subthalamic nucleus (STN) recordings in patients with Parkinson’s disease. This activity is thought to contribute to the gait symptoms of Parkinson’s disease.^35–40^ Since the BG and cerebellum are connected in a functional loop, oscillatory activity in the vermis is likely to be related to oscillatory activity in the STN.^44–45^ Oscillatory activity measurements of low beta and high beta in rats has produced different results from humans. Patients with FOG experienced increased low beta activity during walking in the STN, with no significant increases in the high beta band.^42^ 6-OHDA rats, however, exhibited increased high beta activity in the STN^52^ and in the substantia nigra pars reticulata (SNr)^53^ rats during rest and active gait on a treadmill, respectively. Difference between low beta and high beta activity during forced and volitional gait may give insight into the mechanism behind gait impairment selectivity in Parkinson’s disease. CBLv activity was increased during walking in both beta bands which may be the result of the integration of other cortical regions besides the BG.

Gamma activity is also increased in the BG in Parkinson’s disease. Increases in gamma activity is postulated as a compensatory mechanism for the increase in beta activity but cannot fully compensate due to the inhibitory effect of beta^48^, which could explain the gait improvements in 6-OHDA rats at 21-days. Changes in cerebellar activity from 14- to 21-days may be a response to the 10% increase in DA loss between the timepoints.

Cerebellar activity 28-days post-lesion may be attributed to DA loss. DA loss remained consistent from 21- to 28-days. As the CBLv receives inputs from both hemispheres of BG^45^, this can be attributed to compensation from the non-lesioned side of the brain. Although ~94% DA loss was measured on the lesioned side of the brain, total DA loss from both hemispheres was only about 50%. The STN is hyperactive in Parkinson’s disease^47^ but may function normally on the non-lesioned side of the brain in unilateral 6-OHDA rats. To our knowledge, STN activity in unilaterally lesioned 6-OHDA rats has only been measured ipsilateral to the lesion.^18,46^

CBLv activity decreased during freezing. These changes were measured in the alpha band of lobule VIa and the theta, alpha, high beta, low gamma, and high gamma frequency bands in lobules VIb and VIc at 21-days. Similar activity measured in lobules VIb and VIc supports the correct placement of the μECoG arrays, because prior research has shown that these lobules project to the same brainstem nuclei.^50^ Studies measuring oscillatory activity during walking and standing/rest, have shown increased activity in the high frequency bands (high beta and gamma) and decreased activity in the low frequency (theta, alpha, and low beta) bands during walking compared to standing in the SNr of 6-OHDA rats compared to non-lesioned controls^8,53^,. In humans with FOG, both low beta and high beta power was reduced during standing compared to sitting in the STN.^42^ Our results showed a similar trend in the high frequencies, however, theta and alpha band activity decreased during freezing indicating different oscillatory mechanisms from rest or standing.

Cerebellar theta oscillations are associated with the intermittent control of continuous movements while alpha oscillations are associated with attention and processing.^41^ Together, suppression of these bands may result in the inability to execute motor programs smoothly^49^ resulting in shuffled gait and the inability to re-initiate forward gait. Differences in oscillatory trends between the CBLv and the STN may be an indicator of decoupling between the two regions.^43^ It is unclear from our measures of CBLv oscillatory activity if freezing is truly an indicator of FOG in the 6-OHDA model of Parkinson’s disease. Future research measuring activity before, during, and after freezing in both the vermis and STN concurrently, as well as recording from healthy rats when standing, would provide further insight. These deficits were also improved at 28-days supporting our hypothesis of compensation from the non-lesioned cerebral hemisphere. Additional compensatory mechanisms at play in the lesioned side must also be considered including increased DA release by remaining dopaminergic (DAergic) neurons and DA receptor expression in the striatum.^54^

Research on the CBLv electrophysiological contributions to gait in Parkinson’s disease is lacking. Our results suggest that DAergic cell loss may contribute to cerebellar dysfunction through its disynaptic projections to the BG. The CBLv is hyperactive during walking and reduced during freezing. We believe the mitigation of abnormal activity at 28-days was due to compensation from the non-lesioned side of the brain, which has connections to the CBLv. Further research is needed to determine if the changes measured here are truly indicative of FOG in persons with Parkinson’s disease. Identifying changes in CBLv oscillatory activity immediately before freezing from lobules VIb and VIc may provide a biomarker that can be used in closed-loop DBS to treat FOG. Rat models that present with bilateral dopamine loss may be more indicative of changes in cerebellar activity in response to DAergic cell loss.

## Abbreviations

FOG: Freezing of gait,
SNc: substantia nigra pars compacta,
DA: dopamine,
6-OHDA: 6-hydroxydopamine,
MFB: medial forebrain bundle,
CBLv: cerebellar vermis,
PPN: pedunculopontine nucleus,
BG: basal ganglia,
μECoG: micro-electrocorticography,
STN: subthalamic nucleus,
SNr: substantia nigra pars reticulata,
DAergic: dopaminergic,
FN: fastigial nucleus,
LFP: local field potential,
TH: anti-tyrosine hydroxylase,
DBS: deep brain stimulation.

## References

1 Takakusaki K, Takahashi M, Noguchi T, Chibe R. Neurophysiological mechanisms of gait disturbance in advanced Parkinson’s disease patients. Neurology and Clinical Neuroscience. 2022; 1–17.

2 Bernheimer H, Birkmayer W, Hornykiewicz O, Jellinger K, Seitelberger F. Brain dopamine and the syndromes of Parkinson and Huntington. Clinical, morphological and neurochemical correlations. Journal of the Neurological Sciences. 1973; 20: 415–455.

3 Kish S, Shannak K, Hornykiewicz O. Uneven pattern of dopamine loss in the striatum of patients with idiopathic Parkinson’s disease. Pathophysiologic and clinical implications. New England Journal of Medicine. 1988; 318: 876–880.

4 Morrish PK, Sawle GV, Brooks DJ. Clinical and [18F] dopa PET findings in early Parkinson’s disease. Journal of Neurology, Neurosurgery, and Psychiatry. 1995; 59: 597–600.

5 Garcia-Rill E, Houser CR, Skinner RD, Smith W, and Woodward DJ. Locomotion-inducing sites in the vicinity of the pedunculopontine nucleus. Brain Research Bulletin 1987; 18: 731–738.

6 Chuang C, Su H, Cheng F, Hsu S, Chuang C, Liu C. Quantitative evaluation of motor function before and after engraftment of dopaminergic neurons in a rat model of Parkinson’s disease. Journal of Biomedical Science. 2010; 17: 9.

7 Nutt JG, Bloem BR, Giladi N, Hallet M, Horak FB, Nieuwboer A. Freezing of gait: moving forward on a mysterious clinical phenomenon. The Lancet Neurology. 2011; 10(8): 734–744.

8 Wenger N, Vogt A, Skrobot M, et al. Rodent models for gait network disorders in Parkinson’s disease- a translational perspective. Experimental Neurology. 2022; 352.

9 Amoozegar S, Pooyan M, Roghani M. Identification of effective features of LFP signal for making closed-loop deep brain stimulation in parkinsonian rats. Medical and Biological Engineering and Computing. 2022; 60(1): 135–149.

10 Zhou M, Zhang W, Chang J, et al. Gait analysis in three different 6-hydroxydopamine rat models of Parkinson’s disease. Neuroscience Letters. 2015; 584: 184–189.

11 Simola N, Morelli M, Carta AR. The 6-hydroxydopamine model of Parkinson’s disease. Neurotoxicity Research. 2007; 11: 151–167.

12 Wu T, Hallett M. A functional MRI study of automatic movements in patients with Parkinson’s disease. Brain. 2005; 128(10): 2250–2259.

13 Hanakawa T, Katsumi Y, Fukuyama H, et al. Mechanisms underlying gait disturbance in Parkinson’s disease: a single photon emission computed tomography study. Brain. 1999; 122(7): 1271–1282.

14 Maiti B, Rawson KS, Tanenbaum AB, et al. Functional connectivity of vermis correlates with future gait impairments in Parkinson’s disease. Movement Disorders. 2021; 36(11): 2259–2568.

15 Yu H, Sternad D, Corcos DM, Vaillancourt DE. Role of hyperactive cerebellum and motor cortex in Parkinson’s disease. NeuroImage. 2007; 35(1): 222–233.

16 Rascol O, Sabatini U, Fabre N, et al. The ipsilateral cerebellar hemisphere is overactive during hand movements in akinetic parkinsonian patients. Brain. 1997; 120(1): 103–110.

17 Caligiore D, Helmich R, Hallet M, et al. Parkinson’s disease as a system-level disorder. npj parkinson’s disease. 2016; 2(1): 16025.

18 Sutton AC, O’Connor KA, Pilitsis JG, Shin DS. Stimulation of the subthalamic nucleus engages the cerebellum for motor function in parkinsonian rats. Brain Structure and Function. 2015; 220(6): 3595–3609.

19 Bosch TJ, Groth C, Eldridge TA, Gnimipieba EZ, Baugh LA, Singh A. Altered cerebellar oscillations in Parkinson’s disease patients during cognitive and motor tasks. Neuroscience. 2021; 475: 185–196.

20 Buckner RL. The cerebellum and cognitive function: 25 years of insight from anatomy and neuroimaging. Neuron. 2013; 80(3): 807–815.

21 Chiang CH, Wang C, Barth K, et al. Flexible, high-resolution thin-film electrodes for human and animal neural research. Journal of Neural Engineering. 2021; 18(4).

22 Paxinos G, Watson C. The rat brain, in stereotaxic coordinates. San Diego Academic Press, 1997.

23 McConnell GC, So RQ, Hilliard JD, Lopomo P, Grill WM. Effective deep brain stimulation suppresses low-frequency network oscillations in the basal ganglia by regularizing neural firing patterns. The Journal of Neuroscience. 2012; 32(45): 15657–15668.

24 DeAngelo VM, Hilliard JD, McConnell MC. Dopaminergic but not cholinergic neurodegeneration is correlated with gait disturbances in PINK1 knockout rats. Behavioural Brain Research. 2022; 417.

25 Tremblay SA, Chapman CA, Courtmanche R. State-dependent entrainment of prefrontal cortex local field potential activity following patterned stimulation of the cerebellar vermis. Frontiers in Systems Neuroscience. 2019; 13.

26 Abràmoff MD, Magalhães PJ, Ram SJ. Image Processing with ImageJ. Biophotonics International. 2004; 11(7): 36–42.

27 Ferreira T, Rasband W. The ImageJ User Guide – Version1.44, http://imagej.nih.gov/ij/docs/user-guide.Parkinson’sdiseasef, February 2011.

28 Koziol LF, Budding D, Andreasen N, et al. Consensus paper: the cerebellum’s role in movement and cognition. Cerebellum. 2014; 13(1): 151–177.

29 Stoodley CJ, Schmahmann JD. Chapter 4: functional topography of the human cerebellum. Handbook of Clinical Neurology. 2018; 154: 59–70.

30 Fujita H, Kodama T, Lac SD. Modular output circuits of the fastigial nucleus for diverse motor and nonmotor functions of the cerebellar vermis. eLife. 2020; 9: e58613.

31 Hsieh T, Chen J, Chen L, Chiang P, Lee H. Time-course gait analysis of hemiparkinsonian rats following 6-hydroxydopamine lesion. Behavioural Brain Research. 2011; 222(1): 1–9.

32 Iansek R, Danoudis M. Freezing of gait in Parkinson’s disease: its pathophysiology and pragmatic approaches to management. Movement Disorders Clinical Practice. 2016; 4(3): 290–297.

33 Kwok JYY, Smith R, Chan LML, et al. Managing freezing of gait in Parkinson’s disease: a systematic review and network meta-analysis. Journal of Neurology. 2022; 269(6): 3310–3324.

34 Caligiore D, Pezzulo G, Baldassarre G, et al. Consensus paper: towards a system-level view of cerebellar function: the interplay between cerebellum, basal ganglia, and cortex. The Cerebellum. 2017; 16: 203–229.

35 Alegre M, Alonso-Frech F, Rodriguez-Oroz MC, et al. Movement-related changes in oscillatory activity in the human subthalamic nucleus: ipsilateral vs. contralateral movements. European Journal of Neuroscience. 2005; 22: 2315–2324.

36 Brown P. Oscillatory nature of human basal ganglia activity: relationship to the pathophysiology of Parkinson’s disease. Movement Disorders. 2003; 18: 357–363.

37 Brown P, Oliviero A, Mazzone P, Insola A, Tonali P, DiLazzaro V. Dopamine dependency of oscillations between subthalamic nucleus and pallidum in Parkinson’s disease. The Journal of Neuroscience. 2001; 21(3): 1033–1038.

38 Cassidy M, Mazzone P, Oliviero A, et al. Movement-related changes in synchronization in the human basal ganglia. Brain. 2002; 125(6): 1235–1246.

39 Foffani G, Bianchi AM, Baselli G, Priori A. Movement-related frequency modulation of beta oscillatory activity in the human subthalamic nucleus. Journal of Physiology. 2005; 568(2): 699–711.

40 Kuhn A, Williams D, Kupsch A, et al. Event-related beta desynchronization in human subthalamic nucleus correlates with motor performance. Brain. 2004; 127(4): 735–746.

41 De Zeeuw CI, Hoebeek F, Schonewille M. Causes and consequences of oscillations in the cerebellar cortex. Neuron. 2008; 58(5): 655–658.

42 Singh A, Plate A, Kammermeier S, Mehrkens J, Ilmberger J, Botzel K. Freezing of gait-related oscillatory activity in the human subthalamic nucleus. Basal Ganglia. 2013; 3(1): 25–32.

43 Pozzi NG, Canessa A, Palmisano C, et al. Freezing of gait in Parkinson’s disease reflects a sudden derangement of locomotor network dynamics. Brain. 2019; 142(7): 2037–2050.

44 Wu T, Hallet M. The cerebellum in Parkinson’s disease. Brain. 2013; 136(3): 696–709.

45 Jwair S, Coulon P, Ruigrok TJH. Disynaptic subthalamic input to the posterior cerebellum in rat. Frontiers in Neuroanatomy. 2017; 11: 1–11.

46 Bouali-Benazzouz ZN, Gao D, Benabid AL, Benazzouz A. Intrasubthalamic injection of 6-hydroxydopamine induces changes in the firing rate and pattern of subthalamic nucleus neurons in the rat. Synapse. 2001; 40(2): 145–153.

47 Baunez C, Gubellini P. Chapter 12- Effects of GPi and STN inactivation on physiological, motor, cognitive and motivational processes in animal models of Parkinson’s disease. Progress in Brain Research. 2010; 183: 235–258.

48 Florin E, Erasmi R, Reck C, et al. Does increased gamma activity in patients suffering from Parkinson’s disease counteract the movement inhibiting beta activity? Neuroscience. 2013; 237: 42–50.

49 Thevathasan W, Pogosyan A, Hyam JA, et al. Alpha oscillations in the pedunculopontine nucleus correlate with gait performance in parkinsonism. Brain. 2012; 135: 148–160.

50 Price JB, Rusheen AE, Barath AS, et al. Clinical applications of neurochemical and electrophysiological measurements for closed-loop neurostimulation. Journal of Neurosurgery. 2020; 49(1): e6.

51 Rondi-Reig L, Paradis AL, Lefort JM, Bayan BM, Tobin C. How the cerebellum may monitor information for spatial representation. Frontiers in Systems Neuroscience. 2014; 8: 205.

52 Mottaghi S, Kohl S, Biemann D, et al. Bilateral intracranial beta activity during forced and spontaneous movements in a 6-OHDA hemi-Parkinson’s disease rat model. Frontiers in Neuroscience. 2021; 15: 700672.

53 Avila I, Parr-Brownlie LC, Brazhnik E, Castaneda E, Bergstrom DA, Walters JR. Beta frequency synchronization in basal ganglia output during rest and walk in a hemiparkinsonian rat. Experimental Neurology. 2010; 221(2): 307–319.

54 Smith Y, Villalba R. Striatal and extrastriatal dopamine in the basal ganglia: An overview of its anatomical organization and normal and Parkinsonian brains. Movement Disorders. 2008: 23(S3): S534–S547.

