## Supplementary Figures for "Cerebellar Activity in Hemi-Parkinsonian Rats during Volitional Gait and Freezing"

### Supplementary Material

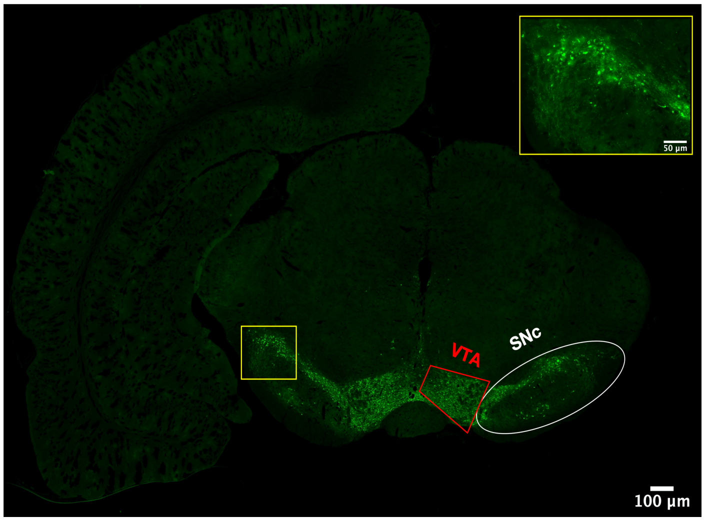


**Supplementary Figure 1: Sample brain sections.** Anti-TH staining to measure dopaminergic neurons in the substantia nigra pars compacta (SNc). The ventral tegmental area (VTA) was spared in 6-OHDA rats by desipramine injection prior to 6-OHDA infusion.

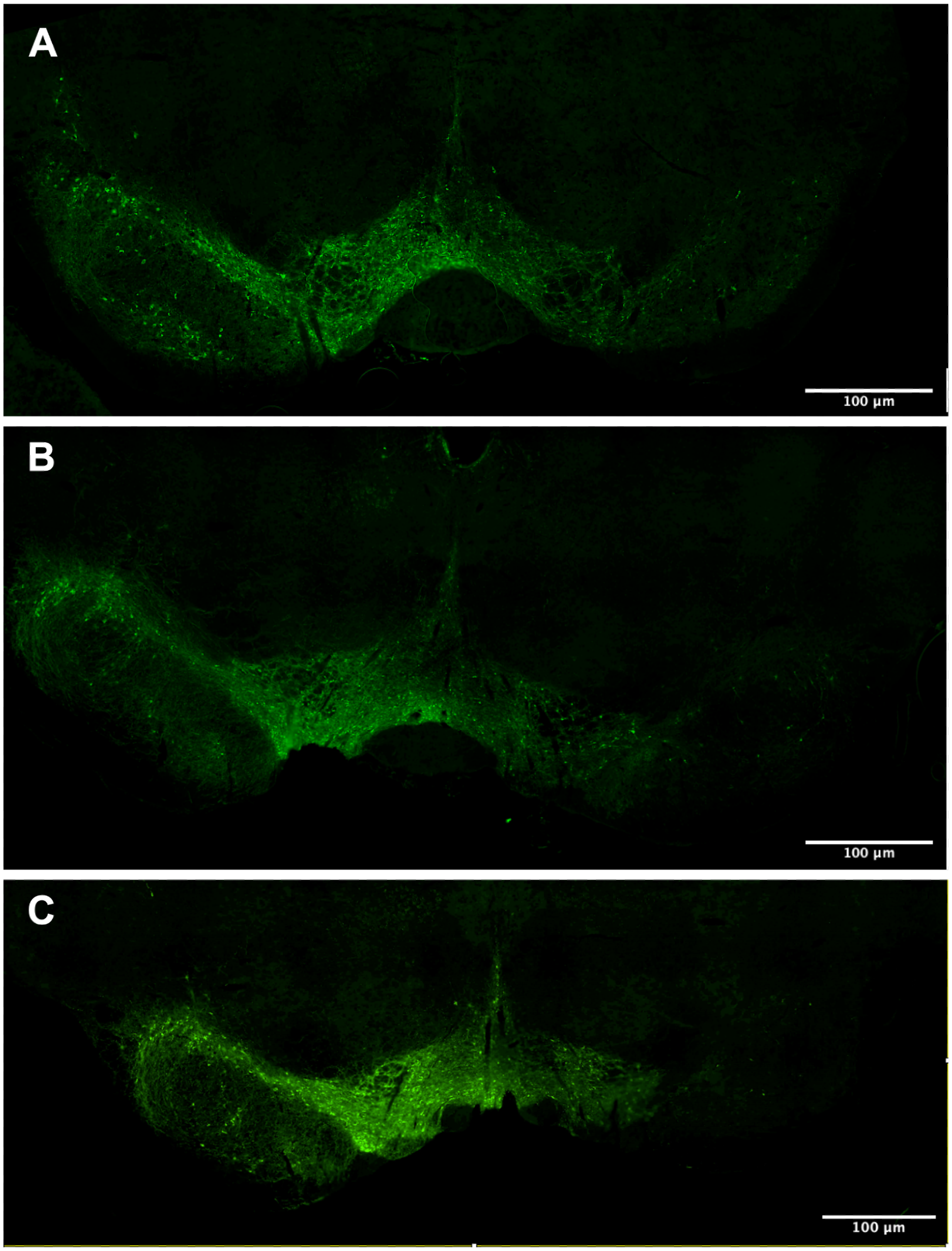


**Supplementary Figure 2: Tyrosine hydroxylase immunohistochemistry.** Dopaminergic TH+ cell loss in the SNc at (A) 14-, (B) 21-, and (C) 28- days after infusion of 6-OHDA in the medial forebrain bundle (MFB). Left hemisphere is non-lesioned side and right hemisphere is lesioned side for all images.
